# Phase Separation of Intracellular Proteins Influences Synaptic Bistability

**DOI:** 10.64898/2026.10.24.761593

**Authors:** Vidhi Saxena, Anagha Anil Totade, Vikas Pandey, Gaurav Chauhan

**Affiliations:** Department of Chemical Engineering, Indian Institute of Technology Indore, Indore, Madhya Pradesh, India 453552; Premium Research Institute for Human Metaverse Medicine (PRIMe), The University of Osaka, Osaka, Japan 565-0871

## Abstract

Synaptic bistability at the post synaptic density has been proposed to be the result of the nucleation and evaporation of receptor clusters. The exact biophysical mechanism underlying these inter-receptor attractions that can trigger nucleation remains to be elucidated. Emerging evidence demonstrates that numerous postsynaptic density submembrane proteins, that interact with various membrane receptors, can undergo phase separation, necessitating a revised conceptual model. By employing Langevin dynamics simulations of coarse-grained scaffold proteins and interacting membrane receptors, we study how phase transitions of scaffold proteins alter receptor clustering at the membrane. In our model, scaffold macromolecules can undergo phase separation and interact with receptor molecules at the model membranes. We observed that the presence of scaffold polymers across the membrane creates an attractive interaction between otherwise purely repulsive receptors. By systematically varying the number of receptors at the membrane, we dissected the coupling between receptor clustering and the phase behavior of scaffold macromolecules. We show that the propensity for receptor clustering is enhanced by the condensation of scaffold proteins. We then used two different scaffolds and systematically varied the interaction between them, to change the spatial organization of two condensed phases w.r.t. one another. Interestingly, we observed that receptor clustering can be reduced by co-phase separation of scaffold macromolecules, indicating the role of competing interactions in receptor clustering. Our work advances understanding of receptor-mediated signaling and phase separation, providing insights into cellular compartmentalization and biochemical regulation at the synapse.

## I. INTRODUCTION

In neuronal signaling, neurons transmit information via synaptic transmission across specialized intercellular junctions known as synapses.[1]. The synapse enables the transmission and alteration of information within neural circuits by converting electrical impulses into chemical signals.[2]. This process is governed by stringently regulated sequence of events such as synaptic vesicle docking, neurotransmitter release, membrane receptor activation, and signal transduction. These are all carefully controlled processes that are controlled by different molecular machinery in the presynaptic and postsynaptic compartments.[3].

Receptor clustering on the plasma membrane is a well-observed neuronal signaling pathway and is also important in other biological contexts.[4, 5] The receptors at the synapse are known to show bistability. By switching between two distinct metastable states containing different numbers of localized receptors, this bistability allows the synapse to maintain long-term stability of synaptic efficacy despite the rapid turnover and short dwell time of individual receptors.[4–6] The existing model for observing bistability at the synapse was proposed to be a result of vapor-liquid transition of the receptor molecules.[6] In this framework, it was essential to have attractive interaction between the receptors to trigger clustering and observe bistability between a low density vapor state and a high density liquid state, depending upon the receptor flux. What remains unclear are the origins of this receptor-receptor interaction and whether there is any role of the submembrane proteins[7] in cluster stabilization.Phase separation of submembrane post synaptic proteins has also been shown to play a role in receptor clustering at the membrane,[4] although a mechanistic understanding is still not clear.

Phase separation of proteins is emerging to be an important mechanism in neuronal signaling. Phase separation is the process by which biomacromolecules, such as proteins, separate from a homogeneous aqueous mixture to form distinct compartments within cells [8, 9]. These compartments are often called membraneless organelles and also referred to as biomolecular condensates [10, 11]. The phase behavior of these biomolecular condensates can be modulated by a range of physical driving forces, including macromolecular crowding and active motility [12, 13]. In neurons, the active zone, endocytic sites, and postsynaptic density are just a few of the synapse types that have been identified as membraneless compartments in recent years [14]. These biomolecules autonomously aggregate into a compact dense phase devoid of any enclosing membrane upon attaining the saturation concentration in a bulk solution [15]. Synaptic transmission and synaptic plasticity are supported by the creation of several sub-compartments, which enable different synaptic molecules/components to be localized and enriched in particular subsynaptic areas [16]. A large body of research has identified a family of scaffold proteins, including RIMs, RIM-BPs, ELKS, liprins, and Munc13s as crucial molecular organizers of active zones that connect voltage-gated Ca^2+^ channels (VGCCs) to the synaptic vesicle fusion machinery [17, 18].

Phase separation of proteins such as postsynaptic density protein 95 (PSD-95) and synapsin-1 is considered important for neuronal signaling. Various receptors, such as AMPA receptors (AMPARs) and NMDA receptors (NMDARs), have been shown to cluster in response to PSD-95 [19]. Scaffolds such as PSD-95, Homer, Shank, and synapsin-1—in both single-phase and multiphase states [20, 21]—organize receptors, including NMDA, AMPA, GABA, and TrkB, in pre- and postsynaptic regions and influence synaptic signaling [22]. However, there is still a lack of mechanistic understanding of how the phase separation of submembrane proteins affects receptor clustering at the membrane.

In this work, we investigate how the phase behaviour of a submembrane scaffold protein governs receptor clustering at the plasma membrane. Specifically, we address two questions: (i) how receptor clustering depends on the scaffold-polymer phase behaviour, and (ii) how the composition and organization of two scaffold macro-molecule condensates modulate receptor clustering. Together, our findings establish how scaffold-driven phase transitions dictate the assembly of signaling-responsive receptor bistability. This provides a quantitative, mechanistic basis for understanding how spatial organization enables cells to tune signal initiation, amplification, and specificity.

## II. METHODS

### A. Coarse-grained modeling of the Protein-Receptor system

We investigated the effect of scaffold-protein condensate formation and dissolution on receptor clustering, using an implicit-solvent Langevin dynamics simulation. We modeled proteins as coarse-grained Kremer-Grest polymers with beads of polymers interacting via the Lennard-Jones potential to examine how scaffold-protein phase behavior affects receptor clustering at the plasma membrane (Fig. 1).

**FIG. 1.**
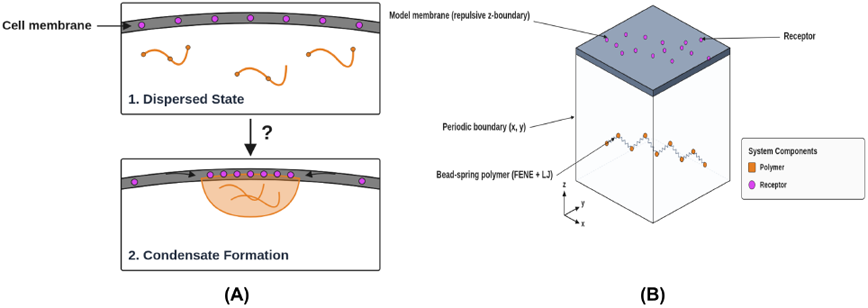
Coarse-grained model of scaffold protein and receptor system. (A) Schematic representation of scaffold proteins and their associated receptors as a bead spring polymer. (B) The system contains a single polymer in the simulation box, with its associated receptor confined to the upper slab. The upper slab has a repulsive z boundary to prevent receptors from entering the lower region and polymers from entering the upper region.

We considered both phase-separated and mixed states over a range of receptor concentrations. The scaffold protein was modeled as a bead-spring polymer comprising *N* =25 monomers per chain, whereas the receptor was modeled as a monomer with varying numbers of *R* = 50, 100, 150, 200, 250, and 300 (Fig. 1B). We considered two cases: in the first case, scaffold polymer (P) beads were modeled as bead-spring polymers with non-connected beads interacting via the Lennard-Jones (LJ) potential, as shown in equation (1).

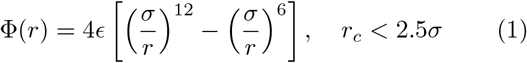

with *ϵ* = 0.5*k*_*B*_*T/σ*. (Ref. [23]). We used a finite extensible non-linear elastic (FENE) potential

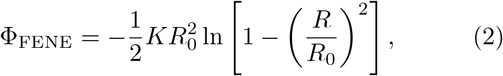

between polymer’s adjacent monomer beads with *K* = 10 and *R*_0_ = 2.0. In the second case, we considered polymers which did not phase separate by using the WCA potential between the monomers, with LJ interactions cut off at *r*_*c*_ = 2^1*/*6^. All simulations were performed using standard Lennard-Jones reduced units, where length is scaled by *σ*, energy by *ϵ*, force constants by *ϵ/σ*^2^, and time by 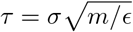.

### B. Coarse-grained modeling of the Membrane

The simulation box was partitioned along the *z*-axis into upper and lower sub-domains separated by a purely repulsive boundary along the *z* dimension (Weeks– Chandler–Andersen potential). This repulsive barrier was parameterized to strictly confine monomeric receptors to the upper compartment and prevent scaffold polymers from traversing from the lower to the upper compartment, thereby establishing a spatially restricted interaction domain. The receptors were treated as particles with LJ interactions between the polymers and the receptors. We considered purely repulsive WCA interactions between the receptors. For the initial set of simulations containing one receptor and one scaffold polymer in the upper and lower box regions respectively, the box dimensions were kept 60 *×* 60 *×* 70.

### C. Coarse-grained modeling of the two Polymer-Receptor system

In the second set of simulations, to study how the multi-component cluster influences receptor clustering, we considered two scaffold polymers (P1 and P2) and two receptors (R1 and R2), each with its respective affinity towards itself and the other polymer. Here, polymer and receptor beads were also modeled using the LJ potential, as in the first case. We considered four cases in which we varied the P1–P2 interaction (*ϵ* = 0.1, 0.25, 0.4, and 0.5) at varying receptor concentrations (number of R1 and R2 = 50, 100, and 204), while keeping the P1-P1 and P2-P2 interaction potential *ϵ* = 0.5*k*_*B*_*T/σ* constant. Table I lists the LJ parameters used in the simulations.

**TABLE I.** Definition of non-bonded interactions between the polymers and receptors of the system.

| Particles | Epsilon | Sigma | Cutoff ($2^{1/6}\sigma$) | Interaction |
| --- | --- | --- | --- | --- |
| P1 – P1 | 0.5 | 1 | 2.5 | Attraction |
| P1 – P2 | 0.1/0.25/0.4/0.5 | 1 | 2.5 | Weak attraction |
| P2 – P2 | 0.5 | 1 | 2.5 | Attraction |
| R1 – P1 | 50 | 1 | 4.0 | Attraction |
| R1 – P2 | 50 | 1 | 1.1225 | Repulsion |
| R2 – P1 | 50 | 1 | 1.1225 | Repulsion |
| R2 – P2 | 50 | 1 | 4.0 | Attraction |
| R1 – R1 | 1.0 | 1 | 1.1225 | Repulsion |
| R1 – R2 | 1.0 | 1 | 1.1225 | Repulsion |
| R2 – R2 | 1.0 | 1 | 1.1225 | Repulsion |

In the second set of simulations with two scaffold polymers and two receptors, the system box dimensions were scaled to 80 *×* 80 *×* 70.

Simulations were performed in the NVT ensemble using the LAMMPS simulation package (23 Jun 2022) [24]. The Langevin equation was integrated forward in time using the velocity-Verlet algorithm. The integration time step was chosen to be 0.005*τ*, where *τ* is the natural unit of time. The damping parameter was set to 1*/τ* . Scaffold polymers and receptor particles were initialized randomly in the box. We then performed Langevin dynamics simulations. Trajectories were recorded at 5 *×* 10^4^ time-step intervals, and each simulation was executed in a total of 2 *×* 10^7^ time steps. In the two-scaffold polymer-receptor system with *ε* = 0.1, we carried out longer simulations by further running for 2 *×* 10^7^ time steps to achieve equilibration. We used the last 200 equilibrated trajectory frames to calculate the fraction of molecules in a cluster. We used OVITO [25] to visualize the simulation trajectories.

## III. RESULTS AND DISCUSSION

Using coarse-grained Langevin dynamics simulations of scaffold proteins that bind to or have affinity with their membrane receptors, we studied the dynamics of receptor clustering in response to the phase separation of scaffold proteins.

### A. Submembrane scaffold protein phase separation enhances the driving force for receptor clustering

We investigated how scaffold protein condensation alters receptor clustering. We studied the system under two conditions: a dispersed phase of the protein (polymer) and a phase-separated two-phase system. Scaffold proteins with pure volume exclusion do not form any dense phase region. We did not observe any clustering up to *R* = 100 (Fig. 2A,B). Beyond this receptor concentration, clustering of receptors is observed due to cooperativity in polymer-mediated interactions between otherwise repulsive receptors (Fig. 2C). We note that the presence of bulk polymers induces an effective interaction between membrane receptors because a single polymer chain can simultaneously bind to multiple receptors on the membrane surface.

**FIG. 2.**
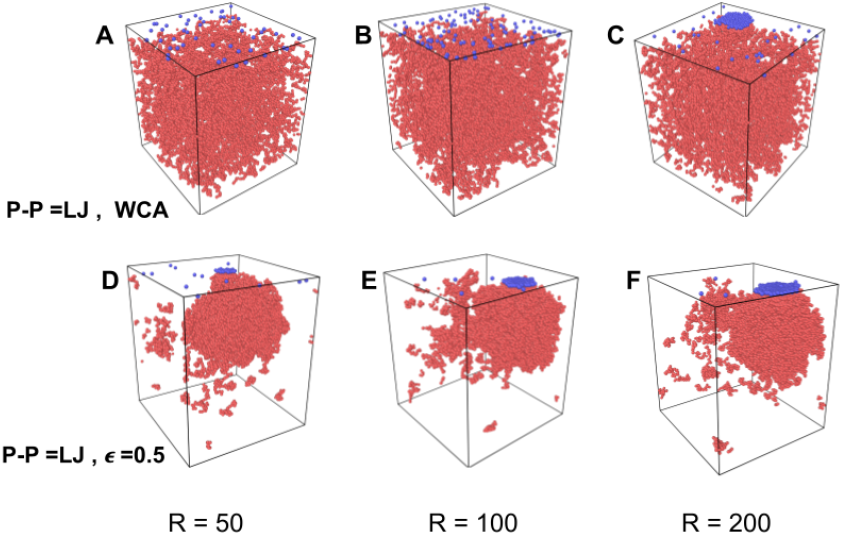
Snapshots of the simulation trajectory showing the spatial organization of the scaffold protein–receptor system. The top row (A–C) corresponds to P–P = WCA, and the bottom row (D–F) corresponds to P–P = LJ, *ϵ* = 0.5. From left to right, the panels show *R* = 50, *R* = 100, and *R* = 200, respectively.

Next, we studied clustering response for weakly attractive scaffold polymers. At weak attractive LJ interaction of *ϵ* = 0.5 among monomers of scaffold proteins (Fig. 2D,E,F; red particles), we observed the formation of a dense phase. Strikingly, under this condition, receptor clustering was observed at much low receptor abundance, at *R* = 50, showing that the driving force for clustering is enhanced in the presence of phase separated scaffolds(Fig. 2D).

To quantify the effect of scaffold condensation on receptor clustering, we evaluated the ensemble-averaged fraction of total receptors belonging to the largest cluster, denoted as *F*_receptor, cluster_, as a function of total receptor count *N* . We compared two pairwise interaction models between receptors (P–P): an attractive Lennard-Jones (LJ) potential with interaction strength *ϵ* = 0.5 *k*_*B*_*T*, and a purely repulsive Weeks–Chandler–Andersen (WCA) potential representing baseline excluded-volume interactions between the monomers of the polymer.

For receptors interacting via the attractive LJ potential, In the low receptor abundance regime (e.g. with *R* = 20), receptors remain predominantly in dispersed states, yielding an ensemble-averaged cluster fraction close to 0. As the system enters the transition region (*R* = 50), a sharp nonlinear increase in the largest-cluster fraction occurs (Fig. 3), indicating the rapid nucleation or cluster formation. Then, beyond *R* = 100, the fraction plateaus, reflecting a saturation regime with a small free monomer concentration. When this attractive force is absent among polymers (P–P = WCA), cluster formation shifts significantly to higher receptor densities and exhibits a distinctly different thermodynamic profile. At low to moderate densities of receptors (*R* = 50, 100, 125), clustering is not observed. The system subsequently undergoes a pronounced density-dependent rise in clustering between *R* = 125 and *R* = 200. This behavior highlights how cooperativity in binding can give rise to receptor clustering at large receptor abundance. Importantly, at the high receptor concentration (where *R* = 300), the cluster fraction is slightly above 0.8, whereas in the attractive case (P–P = *ϵ* 0.5), the cluster fractions are approximately 0.98, demonstrating that phase separation of scaffolds results in a higher fraction of receptors undergoing clustering.

**FIG. 3.**
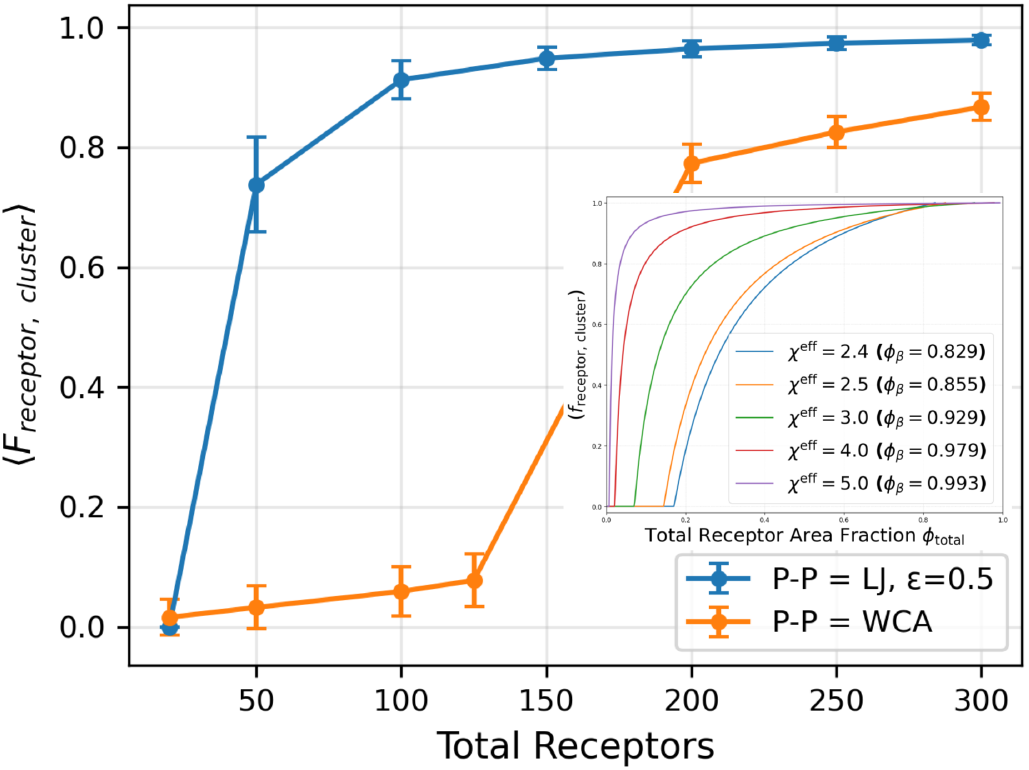
Condensation in the scaffold protein enhances receptor clustering. Under the WCA-only condition (orange), receptors remain largely dispersed at low to intermediate total receptor numbers. A pronounced clustering transition occurs near *R* = 200, after which the clustered fraction increases more gradually with receptor number. In contrast, under the attractive scaffold condition (P–P = LJ, *ϵ* = 0.5; blue), the clustered fraction rises sharply between *R* = 20 and *R* = 50, reaching approximately 0.74 at *R* = 50, and subsequently approaches saturation near unity for *R ≥* 100. Error bars represent the standard error across independent simulation trajectories. The inset shows the predicted clustered receptor fraction, *f*_receptor,cluster_, as a function of the total receptor area fraction, *ϕ*_total_, for different interaction-strength parameter *χ*^*eff*^ (*ϕ*_*P*_ ), and its corresponding clustered phase area fractions, *ϕ*_*β*_. This is obtained by solving the Flory-Huggins/Hildebrand free energy of mixing for a regular solution.

We construct the free energy of mixing of receptors using the Langmuir binding isotherm for polymer site occupancy. We quantify the fraction of polymer-bound receptors as 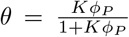, and the fraction of unbound receptors as 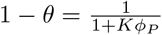. For two neighboring receptor sites *A* and *B* on the membrane, *P* (*A* is bound) = *θ* and *P* (*B* is unbound) = 1 *− θ*. Under mean-field assumptions, the probability of forming a polymer bridge between them is proportional to *θ*(1*− θ*). Therefore, the effective free energy of mixing per lattice site on the membrane can be written as:

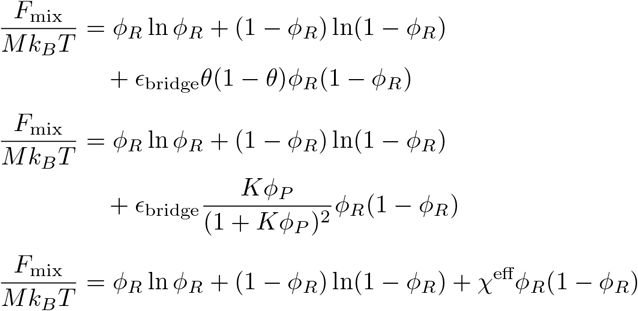

where 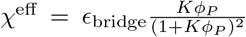 . Here, *F*_mix_ is the total effective free energy of mixing for the receptors on the membrane, *M* is total number of lattice sites on the membrane surface. *ϕ*_*R*_ is the area fraction (or site occupancy fraction) of receptors on the membrane surface.*K* is Equilibrium binding (association) constant between a polymer segment and a receptor, *ϕ*_*P*_ is the bulk volume fraction of the polymer near the membrane interface and *ϵ*_bridge_ is the net free energy gain (in units of *k*_*B*_*T* ) released upon forming a single polymer bridge between two receptors.*χ*^eff^ is the effective receptor–receptor interaction parameter, defined as 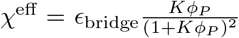 and is a function of the polymer concentration near the membrane interface.

We solve the 2D Flory-Huggins/Hildebrand free energy of mixing numerically to study the fraction of receptors in the cluster as a function of total receptor concentration, for different effective interaction-strength parameter *χ*^*eff*^ (*ϕ*_*P*_ ). Since phase separation increases the polymer concentration near the membrane, the value of *χ*^*eff*^ also increases upon phase separation. We observe that an effective attractive interaction parameter *χ*^*eff*^ (*ϕ*_*P*_ ) can reliably reproduce the simulation results. Therefore, the presence of submembrane proteins only changes the attractive interaction between the receptors and the framework of [6] can be used to understand synaptic bistability. When the submembrane proteins undergo phase separation, it can result in larger concentration of scaffold proteins at the membrane vastly increasing the effective interaction between receptors, as captured by an increased *χ*^*eff*^ (*ϕ*_*P*_ ).

### B. Competing interactions between two scaffold polymers alters receptor clustering response

In the second set of simulations, we investigated how multi-component condensates influences receptor clustering. We considered two polymeric scaffold proteins (P1 and P2) present in the lower region of the simulation box, and two monomeric receptors (R1 and R2) confined to the upper region. Each scaffold protein has strong intraprotein attractive interactions (P1–P1 and P2–P2), while we varied the inter-scaffold protein interaction strengths (P1–P2) (*ϵ* = 0.1, 0.25, 0.4, and 0.5). This led to the formation of different spatial organization of the two condensates, including completely isolated phase-separated condensates of each scaffold protein (*ϵ* = 0.1 and 0.25), condensates that just touch (*ϵ* = 0.4), and co-phase-separation (*ϵ* = 0.5).

Figure 4 shows the clustering response with different spatial organization of the condensates of the two-polymers, at two different receptor concentrations (R1, R2 = 100 and R1, R2 = 204). When the scaffold proteins are phase-separated into two distinct condensates, receptor clustering with their respective affinities is observed (Fig. 4A,B) similar to the isolated condensed phase in Fig. 2. When we increase the interaction strength between polymers P1 and P2 to *ϵ* = 0.1, 0.25, we observe that the condensates are barely touching and we observe receptor clustering similar to the isolated condensate case. However, when we increase the interaction strength between the two polymers P1 and P2 to *ϵ* = 0.40, 0.50, we observe a decrease in receptor clustering at small receptor number. This is because interaction of two different polymers with one another comes at the cost of receptor-polymer interactions and therefore forming a co-phase-separated condensate decreases the tendency for receptor clustering (Fig. 5).

**FIG. 4.**
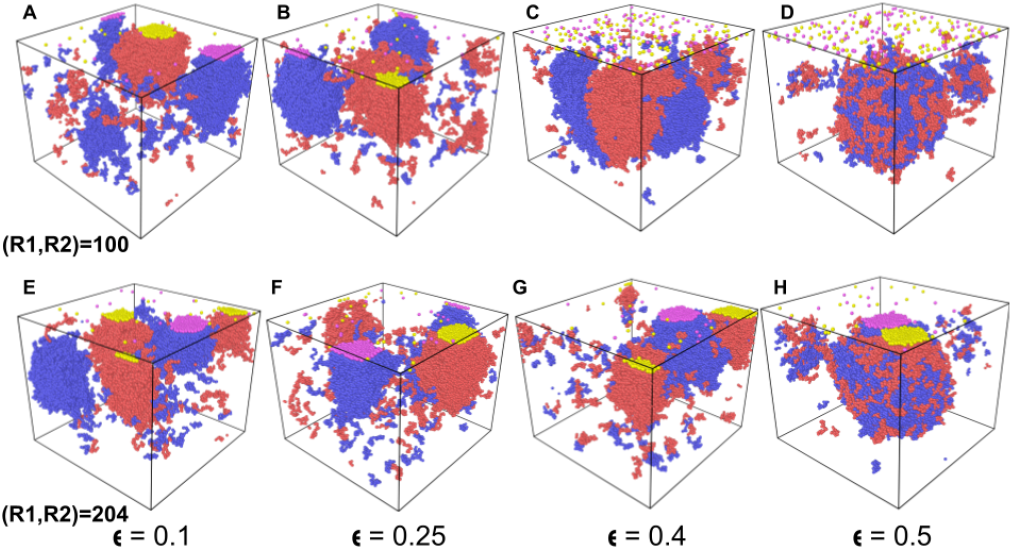
Snapshots of the simulation trajectory showing the spatial organization of the two-scaffold-protein, two-receptor system. The top row (A–D) corresponds to receptor concentration (R1, R2) = 100; the bottom row (E–H) corresponds to receptor concentration (R1, R2) = 204, at different P1–P2 interaction strengths. Top to bottom: A,E at *ϵ*_P1-P2_ = 0.1; B,F at *ϵ*_P1-P2_ = 0.25; C,G at *ϵ*_P1-P2_ = 0.4; and D,H at *ϵ*_P1-P2_ = 0.5.

**FIG. 5.**
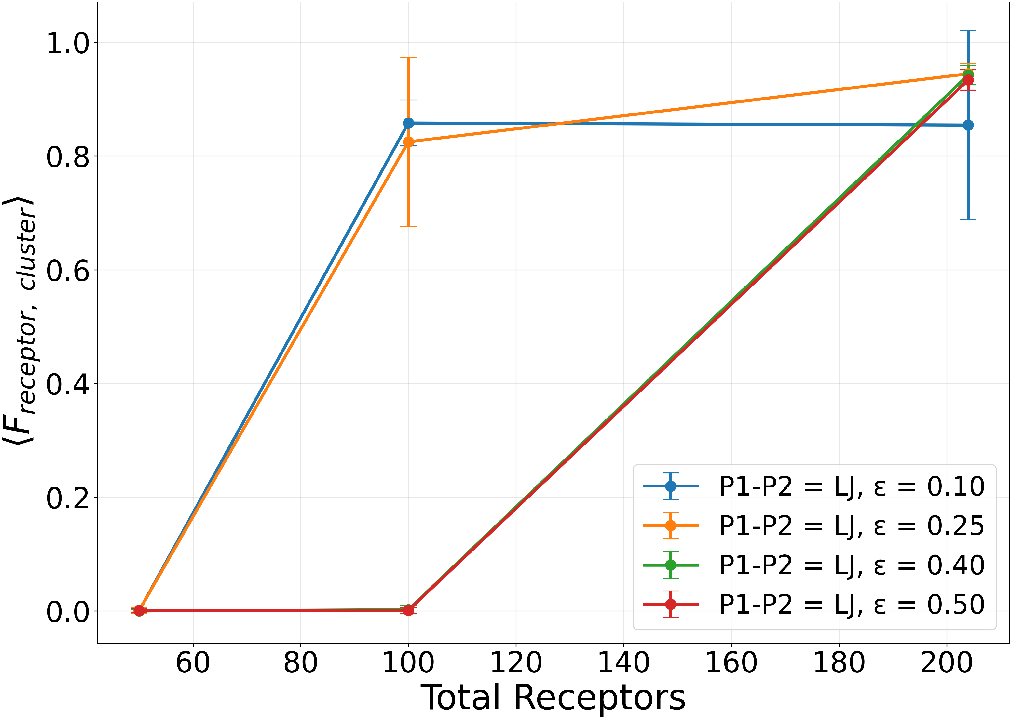
Co-phase separation of proteins suppresses the tendency of receptor clustering. Average fraction of receptors in a cluster plotted as a function of total receptor count across different inter-polymer interaction potential strengths(*ϵ*.The clustering response increases markedly with the number of receptors for attractive LJ interactions. Error bars represent standard errors across simulation trajectories.

There are debates about the function of biomolecular condensates in post synaptic density (PSD). The existing framework of Burlakov *et al*. can explain the observed synaptic bistability of receptors due to liquid-vapor phase transition of receptors. In this work, we show how phase separation of submembrane scaffold proteins in PSD can increase the propensity for receptor clustering at the membrane. This provides a mechanistic explanation for the observed clustering of receptors in the neuronal synapse, and advances our understanding on the role of condensates in neuronal biology.

## IV. DATA AVAILABILITY

The LAMMPS simulation files and analysis code can be obtained from the publicly accessible github repository: https://github.com/SIGNALLING-PAPER/SIGNALLING-PAPER

